# Co-translation drives the assembly of mammalian nuclear multisubunit complexes

**DOI:** 10.1101/419812

**Authors:** Ivanka Kamenova, Pooja Mukherjee, Sascha Conic, Florian Mueller, Farrah El-Saafin, Paul Bardot, Jean-Marie Garnier, Doulaye Dembele, Simona Capponi, H.T. Marc Timmers, Stéphane D. Vincent, László Tora

## Abstract

Cells dedicate significant energy to build proteins often organized in multiprotein assemblies with tightly regulated stoichiometries. As genes encoding proteins assembling in the same multisubunit complexes are dispersed in the genome of eukaryotes, it is unclear how multisubunit complexes assemble. We show that mammalian nuclear transcription complexes (TFIID, TREX-2 and SAGA) composed of a large number of subunits but lacking precise architectural details are built co-translationally. We demonstrate that the dimerization domains and their positions in the interacting subunits determine the co-translational assembly pathway (simultaneous or sequential). Our results indicate that protein translation and complex assembly are linked in building mammalian multisubunit complexes and suggest that co-translational assembly is a general principle in mammalian cells to avoid non-specific interactions and protein aggregation. These findings will significantly advance structural biology by defining endogenous co-translational building blocks in the architecture of multisubunit complexes.

## Introduction

Often proteins do not act alone, instead they function as components of large multisubunit complexes in a cell. To better understand cellular functions, investigating the precise mechanism that guide the formation of these multisubunit assemblies is of key importance. A cell uses hundreds of different protein complexes that vary with respect to their complexity. Some complexes require the association of multiple copies of the same subunit, while others are constituted of many different subunits. The latter group includes many transcription factors and chromatin remodelling complexes (see below). In order to achieve the efficient formation of protein complexes in eukaryotes, the genes coding for all the subunits (dispersed in the eukaryotic genome) have to be transcribed in the nucleus, their corresponding mRNAs transported to the cytoplasm, translated into proteins and the formation of correct interactions amongst the subunits must be orchestrated A polysome is a cluster of ribosomes acting on a single mRNA to translate its information into polypeptides. Appropriate translation-based mechanisms may exist in the cell to regulate the interactions between specific subunits in order to avoid incorrect non-specific interactions or subunit aggregations in the absence of the correct partner. Currently, it is not well understood how functional subunit interactions are regulated in eukaryotic cells. Protein complex formation is often studied *in vitro* using purified subunits, assuming that individually translated subunits assemble stochastically by diffusion, and thus favouring the idea that these multisubunit complexes assemble post-translationally^1^. However, in the crowded environment of an eukaryotic cell such simple diffusion-dependent models may not work, as subunits may engage in non-specific interactions or form aggregates. Recent studies in bacteria demonstrated that co-translational building of a functional protein dimer is more efficient than the post-translational assembly of its individual subunits^2,3^, and also in yeast co-translation has been shown to be an efficient assembly pathway to assemble multiprotein complexes^4-8^. Consequently, two co-translational models have been put forward: i) the “simultaneous” model which suggests that two polysomes in close physical proximity synthesize subunits, which interact while being translated, and ii) the “sequential” model implies that a mature fully translated subunit interacts co-translationally with its polysome-bound nascent interaction partner^9^.

Transcription initiation is a major regulatory step in eukaryotic gene expression. Co-activators establish transcriptionally competent chromatin and promoter environments to allow the formation of the preinitiation complex, comprising RNA polymerase II (Pol II) and general transcription factors (GTFs). Many GTFs and co-activators are multisubunit complexes, in which individual components are organized into functional modules carrying specific activities. In mammalian cells the TFIID GTF nucleates the assembly of the Pol II preinitiation complex on all protein-coding gene promoters [^10,11^ and references therein]. Metazoan TFIID is composed of the TATA-binding protein (TBP) and 13 TBP-associated factors (TAFs) (Fig. 1a). SAGA (Spt Ada Gcn5 Acetyltrasferase) is a multisubunit transcriptional coactivator complex, composed of 19 subunits (including a subset of TAFs), required for the transcription of all active genes in yeast^12^. Moreover, the mammalian Transcription and mRNA Export 2 complex (TREX-2) is composed of five subunits, including the subunit ENY2, which is shared with the SAGA complex^13^.

**Figure 1.**
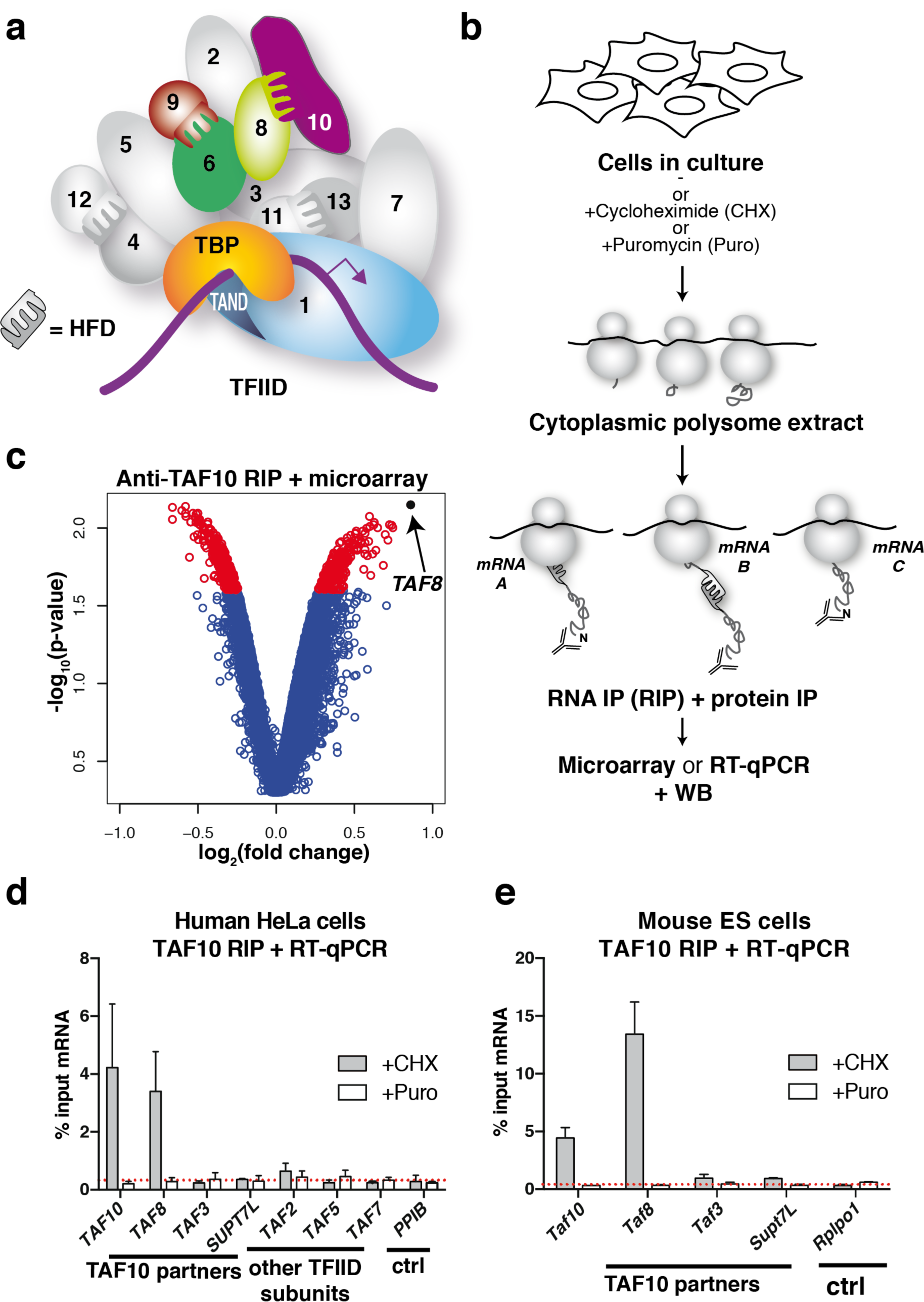
Cotranslational assembly of mammalian TFIID. **(a)** TFIID bound to promoter DNA. TFIID is composed of TBP and 13 TAFs (indicated by numbers). Subunits analysed in this study are highlighted in colour. The histone fold domain (HFD) interactions and the TBP-TAF1 TAND domain interaction are highlighted. **(b)** Schematic representation of polysome RIP assay. **(c)** Endogenous TAF10 was immunoprecipitated from HeLa polysome-containing extract using an antibody targeting the N-terminus of the protein. The enrichment of the precipitated RNAs was assessed globally by microarray. Volcano plot depicting microarray results as log2 of the fold change of IP over a mock IP. A p-value cutoff ≤ 0.025 was applied and corresponding transcripts are in red. *TAF8* transcript is highlighted in black. **(d-e)** RIP-qPCR validation of the microarray results in HeLa (**d**) and mouse ES **(e)** cells. Error bars are ± SD from 3 (HeLa) or 2 (mESC) biological replicates. *Ctrl* = negative control mRNA. *PPIB* and *Rplp0* were used as unrelated control mRNAs.

The majority of TAFs dimerize via their histone-fold domains (HFDs), which are structurally homologous to histone pairs. In TFIID, TAFs form 5 HF pairs (TAF4-12, TAF6-9, TAF8-10, TAF3-10 and TAF11-13) [^10^ and references therein] (Fig. 1a). Importantly, individual HFD-containing TAFs cannot be expressed in a soluble form in bacteria. However, HFD-containing TAFs become soluble when co-expressed with their corresponding specific interaction partner^14^, suggesting that individual HFD-containing TAFs aggregate without their specific partners.

To test how mammalian cells can avoid the aggregation of individual subunits following translation and whether co-translational interactions guide the assembly of transcription complexes, we first investigated pairwise assembly of TFIID subunits between TAF8-TAF10, TAF6-TAF9, TAF1-TBP in polysome-containing mammalian cell extracts. By using a large series of complementary experiments, we show that TAF8-TAF10 and TAF1-TBP assemble co-translationally according to the sequential assembly pathway, while TAF6-TAF9 assemble co-translationally according to the simultaneous model. We also demonstrate that the ENY2 subunit assembles co-translationally with its interaction partner, GANP, in TREX-2, and with ATXN7L3 in the deubiquitination (DUB) module of SAGA. Furthermore, our experiments show that solely the interaction domain (ID) and the position of the ID in the given subunit drives the co-translational assembly in these complexes. Thus, our results open new avenues in the mechanistic understanding of co-translational control of protein complex formation in mammalian cells.

## Results

### TAF10 interacts co-translationally with the polysome-bound nascent TAF8 protein

To test whether HFD-containing TAFs assemble co-translationally, we used a monoclonal antibody against the N-terminus of the HFD-containing TAF10 to immunoprecipitate (IP) endogenous TAF10 from human HeLa cell cytosolic polysome extracts (Fig. 1b). Protein-protein interactions between nascent proteins still associated with translating ribosomes would be revealed by enrichment of mRNAs coding for the interacting partners in the IPs. Global microarray analysis of mRNAs precipitated by the anti-TAF10 RNA IPs (RIPs) revealed enrichment of the *TAF8* mRNA, suggesting that the well-characterized TAF8-10 HFD dimer^15^ forms co-translationally (Fig. 1c). Anti-TAF10 RIP of cytosolic polysome extracts coupled to RT-qPCR validation confirmed our microarray results and showed important enrichment of the *TAF8* mRNA (Fig. 1d). The absence of significant *TAF10* mRNA signal in the microarray experiments was due to weak quality and the high GC-content of the *TAF10* probe sets present on the commercial microarray (data not shown). Nevertheless, RT-qPCR validation also revealed the presence of *TAF10* mRNA in the nascent anti-TAF10 RIP. Importantly, cycloheximide, which “freezes” translating ribosomes on the mRNA^16^, stabilized the TAF10-TAF8-*TAF8* mRNA interactions, while puromycin, which causes release of nascent peptides from ribosomes^17^, resulted in the loss of co-purified mRNA. Endogenous anti-TAF10 RIP-RT-qPCR from polysome extracts prepared from mouse embryonic stem cells (mESCs) gave nearly identical results, which emphasizes the generality of the co-translational pathway for assembly of the mammalian TAF8-TAF10 heterodimer (Fig. 1e; Supplementary Fig. 1). Quantification of the *TAF8* mRNA in the anti-TAF10 RIP normalized to the protein IP efficiency indicated that the enrichment was between 7 and 25%, depending on the cell line and the antibody used. In contrast, to *TAF8*, mRNAs encoding other potential TAF10 dimerization partners, TAF3 and SUPT7L^18^, were not enriched in the RT-qPCR validation experiments, in good agreement with the microarray analysis and indicating the specificity of the co-translational assembly of the TA8-TAF10 heterodimer (Fig. 1d,e). Together these results indicate that TAF10 protein is associated, via the nascent TAF8 protein, with ribosomes which are actively translating *TAF8* mRNA.

### Histone fold domain drives the co-translational assembly of TAF10-TAF8

The fact that TAF8 has its dimerization HFD at an N-terminal position, and that the TAF10 HFD is at the very C-terminus of the protein, allows the direct testing of the sequential assembly model, as TAF8 and TAF10 may be expected to only heterodimerize if the TAF10 protein is fully synthesized and freed from the ribosome. To examine the two assembly models (see Introduction) and to distinguish between the nascent and mature forms of the TAF8 and TAF10 proteins, we added FLAG-, or HA-tags to either N-(to carry out nascent IPs) or C-termini (to carry out mature IPs) of these proteins, respectively. Importantly, exogenous co-expression of N-terminally tagged TAF8 and TAF10 in HeLa cells followed by nascent anti-HA-TAF10 RIP from cytosolic polysome extracts recapitulated the findings obtained with endogenous proteins (Fig. 2a). In contrast, nascent anti-FLAG-TAF8 RIP resulted in high enrichment of its own encoding mRNA, but not that of *TAF10* (Fig. 2b). Immunoprecipitating mature TAF10-HA protein resulted in *TAF8* mRNA, but not *TAF10* mRNA enrichment (Fig. 2c), supporting the sequential co-assembly model, where mature TAF10 released from ribosomes interacts with translating TAF8. Additionally, the mature TAF8-FLAG protein did not bring down any of the tested mRNAs (Fig. 2d). In all cases, protein partners were co-immunoprecipitated successfully (Supplementary Fig. 2). Taken together, these results suggest that mature TAF10 binds to the polysome-bound nascent TAF8 protein, and that the respective N-(in TAF8) and C-terminal (in TAF10) HFDs are driving co-translational dimerization.

**Figure 2.**
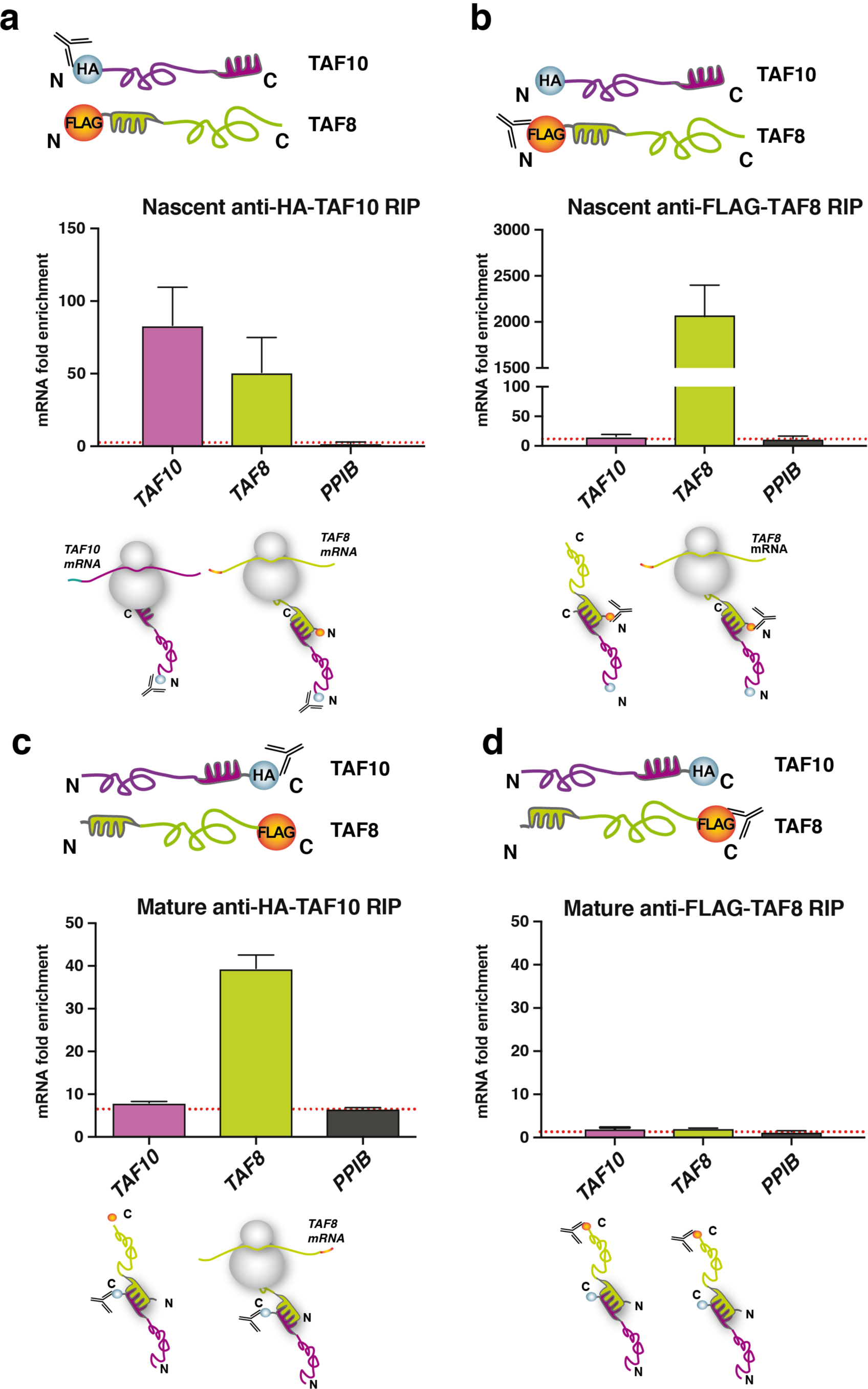
Sequential assembly of TAF10 and TAF8. N-**(a)** and C-terminal **(c)** anti-HA RIP-qPCR of HeLa cell polysome extracts, co-transfected with the corresponding TAF8 and TAF10 expression plasmids (as indicated). N-**(b)** and C-terminal **(d)** anti**-**FLAG RIP-qPCR of HeLa cell polysome extracts, co-transfected with the indicated expression plasmids. *PPIB* **(a -d)** was used as negative control mRNA. In panels (**a-d**) enrichment is expressed as fold change with respect to the mock IP calculated by the formula ΔΔCp [IP/mock] and error bars represent ±S.D. from 2 biological replicates.

To test whether the observed co-translational TAF8-TAF10 assembly is specific to the dimerization of their HFDs, we engineered a mutation disrupting the dimerization ability of the TAF8 HFD (see Methods). Anti-TAF10 RIP from cells co-transfected with *TAF10* cDNA and mutant HFD expressing *TAF8* cDNA (mtTAF8) resulted in a nearly complete loss of the co-precipitated *TAF8* mRNA and TAF8 protein, as compared to the wild type controls (Fig. 3a and Supplementary Fig. 3a), indicating that the dimerization of TAF8 and TAF10 through their HFDs is crucial for co-translational assembly.

**Figure 3.**
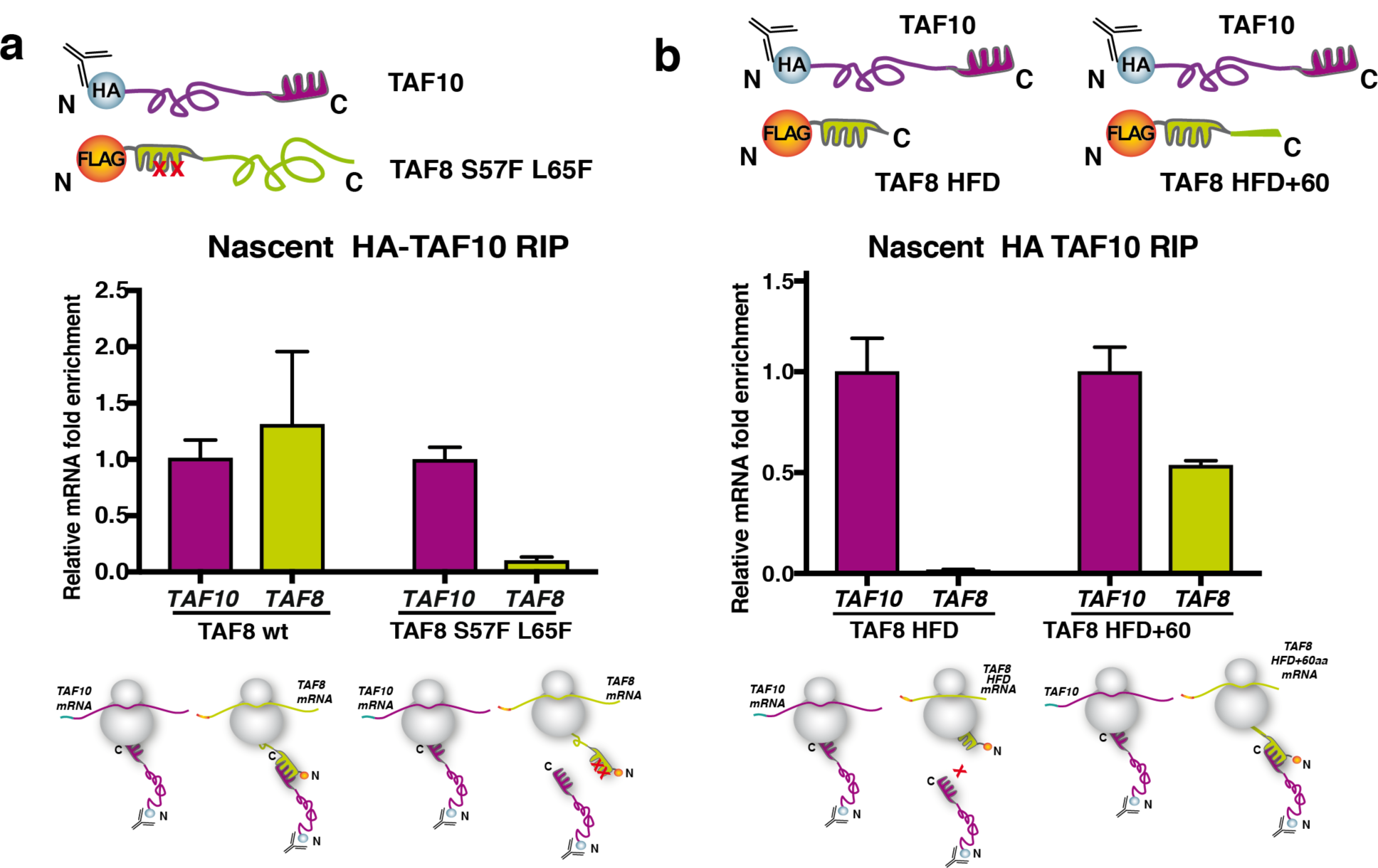
Protein-protein ID drives cotranslational assembly of TAF10 and TAF8. **(a)** anti-HA-TAF10 RIP-qPCR of HeLa cells co-transfected with HA-TAF10 and either wild type FLAG-TAF8 (TAF8 wt) or mutant FLAG-TAF8 (TAF8 S57F L65F). **(b) anti-**HA-TAF10 RIP-qPCR of HeLa cells co-transfected with HA-TAF10 and either a minimal TAF8 HFD or TAF8 HFD extended with 60 amino acids (TAF8 HFD+60aa). In panels (**a-b**) enrichment is expressed as fold change with respect to the mock IP calculated by the formula ΔΔCp [IP/mock] and error bars represent ±S.D. from 2 biological replicates and mRNA enrichment relative to *TAF10* mRNA is represented.

Next, we tested whether the full exposure of the nascent HF interaction domain at the ribosomal exit tunnel would be necessary for co-translational assembly. The ribosome exit tunnel can accommodate up to 60 amino acids [^19^ and references therein]. Thus, we constructed two truncated versions of TAF8: one encoding only the TAF8 HFD that would be partially buried in the ribosome exit tunnel during translation, and a second encoding the TAF8 HFD and an additional 60 amino acids of TAF8 (TAF8 HFD+60) that would allow the appearance of the nascent TAF8 HFD from the ribosomal tunnel. Next a TAF10 expressing plasmid was co-transfected either with TAF8 HFD, or TAF8 HFD+60 expressing plasmids and anti-TAF10 RIPs were carried out. Importantly, our results show that the *TAF8 HFD* mRNA is not enriched in the anti-TAF10 RIP, indicating that the minimal TAF8 HFD protein is released immediately from translating polysomes without co-translational binding to TAF10 protein. On the other hand, the *TAF8 HFD+60* mRNA was enriched in the anti-TAF10 RIP demonstrating that the additional 60 amino acids in the longer TAF8 HFD+60 protein kept the nascent protein anchored in polysomes allowing for co-translational interaction with TAF10 (Fig. 3b and Supplementary Fig. 3b). Together, our results indicate that TAF8-TAF10 co-translational assembly is driven by dimerization with nascent TAF8 protein upon emergence of its entire HFD from actively translating polysomes. Consequently, these results together demonstrate the sequential co-translational assembly pathway where the fully synthesized TAF10 interacts uni-directionally with the nascent TAF8 polypeptide.

### In the absence of TAF10, *Taf8* mRNA and TAF8 protein are prone to degradation, whereas TAF10 protein is stable in *Taf8* knock-out (KO) mESCs

In the sequential assembly pathway, if nascent chains of a subunit cannot co-translationally interact with its partner, it may become prone to misfolding and degradation by the proteasome, but the fully translated partner should stay stable. To test this hypothesis, we used mESCs in which either the endogenous *Taf10*, or *Taf8* genes can be conditionally knocked out^20,21^. By using these mouse ESCs we observed that the deletion of *Taf10* not only ablated *Taf10* mRNA and TAF10 protein levels, but significantly reduced both *Taf8* mRNA and TAF8 protein expression (Fig. 4a,c). These results were also confirmed in *Taf10* KO mouse embryos^20^. In contrast, the deletion of *Taf8*, decreased only its own mRNA and protein levels, without affecting the *Taf10* mRNA expression and TAF10 protein levels (Fig. 4b,d). Furthermore, in both KO mESCs other tested TFIID subunits remained unchanged [^20^ and data not shown]. Together these results further support our findings, indicating that TAF10 interacts co-translationally with TAF8 uni-directionally and when TAF10 is not present TAF8 is prone for degradation. Thus, the nascent TAF8 interaction domain, in the absence of its interaction partner TAF10, may serve as a signal for both protein and mRNA degradation, while TAF10 is stable in the absence of TAF8.

**Figure 4.**
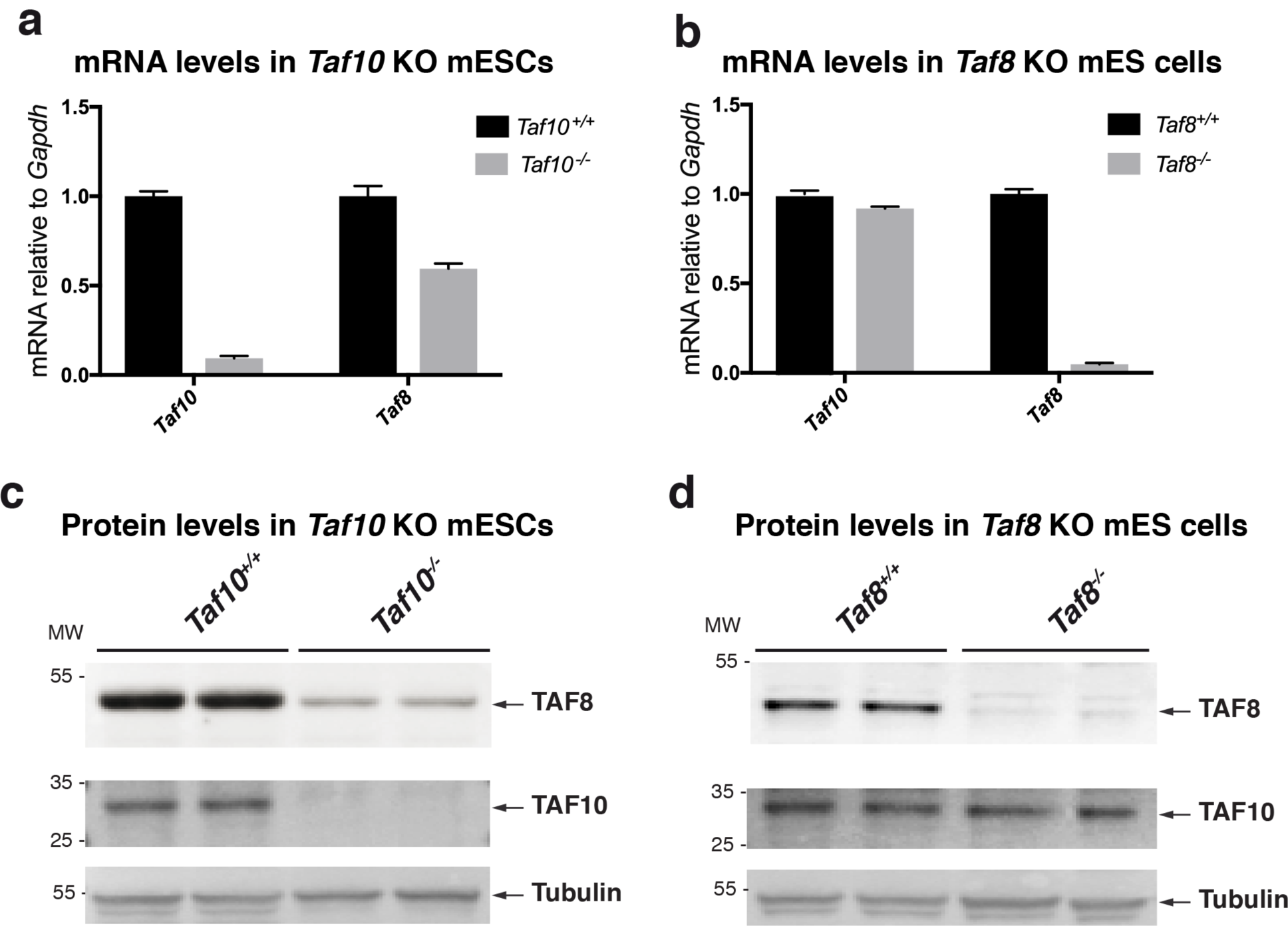
In the absence of TAF10, *Taf8* mRNA and TAF8 protein are prone to degradation in KO mESCs. (**a-b**) RT-qPCR of TAF10 (**a**) and TAF8 (**b**) depleted mESCs. (**c-d**) Western blot analyses from TAF10 (**c**) and TAF8 (**d**) depleted mESCs whole cell extract using anti-TAF8 and anti-TAF10 antibodies. Molecular weight (MW) markers are shown in kDa and an anti-Tubulin was used as a loading control. In panels (**a-b**), mRNA levels were normalized to *Gapdh* mRNA and relative enrichment was calculated using the ΔΔCp method and error bars represent ±S.D. from 3 technical replicates.

### TAF10 protein co-localises with *TAF8* mRNA in the cytoplasm

To visualise the co-localisation of TAF10 protein with *TAF8* mRNA in the cytoplasm, we set out to detect TAF10 protein and *TAF8* mRNA in the cytoplasm of fixed human HeLa cells. To this end we combined protein detection by immunofluorescence (IF) with RNA detection by single molecule inexpensive FISH (smiFISH)^22^. Co-localization of protein and mRNA was then observed by confocal microscopy and quantified. Surprisingly we observed a large difference between the number of endogenous *TAF8* and *TAF10* mRNAs, showing that there are about 4 times less *TAF8* mRNAs than those of *TAF10* (Supplementary Fig. 4a,b). In good agreement with our above endogenous anti-TAF10 RIP results (Fig. 1d,e), these IF-smiFISH experiments showed an about 10% colocalization between TAF8 mRNA and TAF10 protein in the cytoplasm of HeLa cels (Supplementary Fig. 4c). To increase the number of *TAF8* mRNA molecules in the cytoplasm of HeLa cells and to be able to carry out analyses with wild type (wt) and mutant (mt) TAF8 proteins, we carried out IF-smiFISH detections in HeLa cells exogenously expressing TAF8 protein. The IF-smiFISH co-localization experiments in fixed HeLa cells showed significant co-localisation between TAF10 protein and *TAF8* mRNA in the cytoplasm (Fig. 5a,e). Importantly, we lost the co-localisation between *mtTAF8* mRNA (Fig. 3a) and TAF10 protein (Fig. 5b,e). Additionally, TAF8 protein detection by IF and *TAF10* mRNA by smiFISH, showed no significant co-localisation (Fig. 5c,e). Moreover, we could not detect any co-localisation between *CTNNB1* (*catenin beta-1*) mRNA and TAF10 protein (Fig. 5d,e), which further rules out any non-specific co-localisation of TAF10 protein with wt *TAF8* mRNA. Importantly, the statistical analysis of the co-localization enrichment ratio of TAF10 protein-wt *TAF8* mRNA measured in cells was significantly higher compared to all the other conditions tested (Fig. 5e). These imaging experiments demonstrate the physical proximity of TAF10 protein to *TAF8* mRNA in the cytoplasm. Moreover, this proximity is dependent on the ability of the two proteins to interact, lending further support to the sequential assembly model.

**Figure 5.**
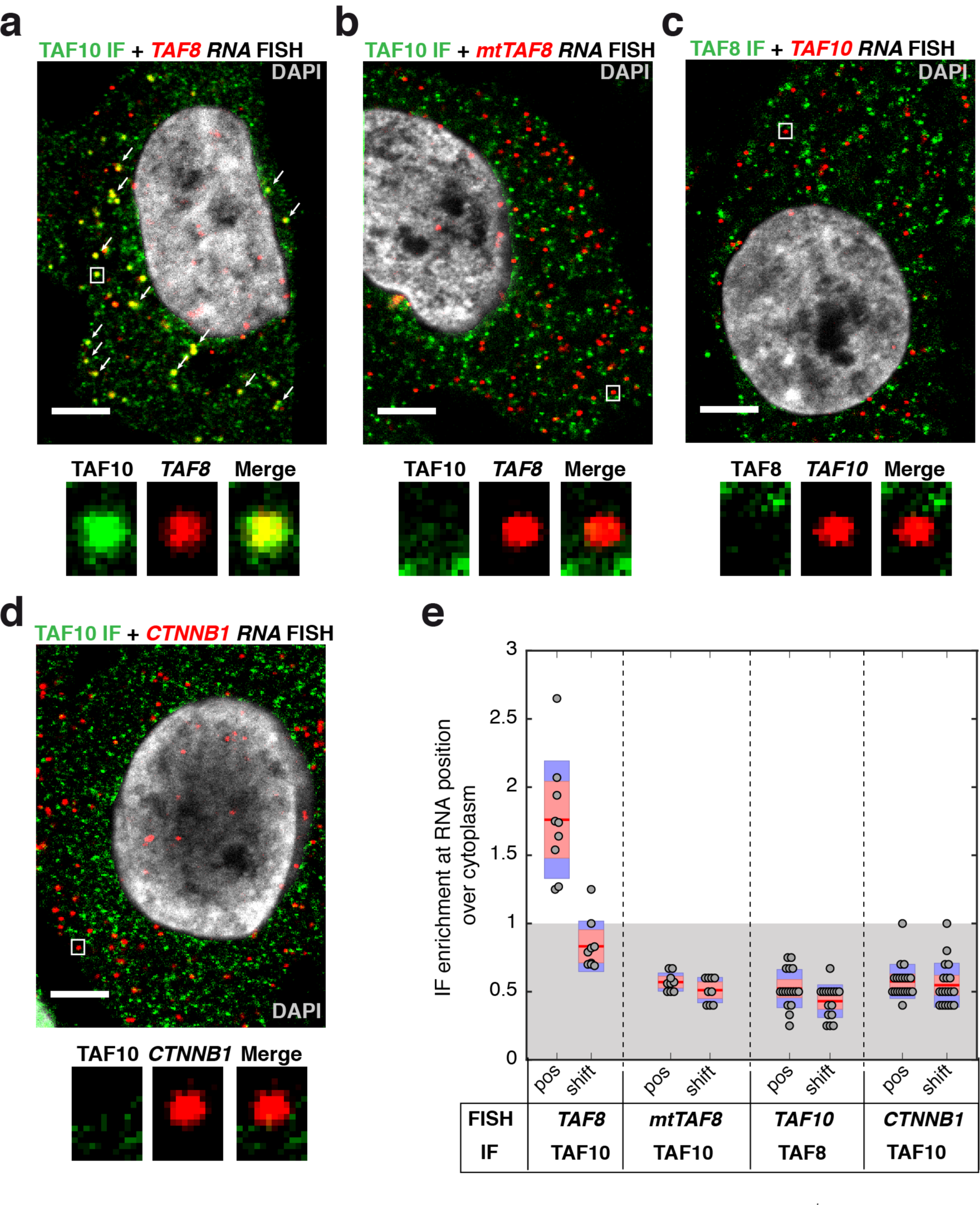
TAF10 protein colocalizes with *TAF8* mRNA in the cytoplasm of HeLa cells. **(a-b)** IF-smiFISH images in HeLa cells expressing either wild type FLAG-TAF8 or mutant (mt) FLAG-TAF8 (TAF8 S57F L65F). Labels: red, Cy3-labelled *TAF8* probes; green, Alexa-488 labelled secondary antibody for TAF10 protein; co-localizing spots are indicated with white arrows. **(c-d)** Representative IF-smiFISH images of endogenous *TAF10* mRNA and TAF8 protein (**c**) and *CTNNB1* mRNA and TAF10 protein (**d**) in HeLa cells. Labels: red, Cy3-tagged *TAF10* FISH probes (**c**) and *CTNNB1* probes (**d**); green, Alexa-488 labelled secondary antibody for TAF8 (**c**) and TAF10 (**d**) protein. A typical cell recorded in each case and after counterstaining with DAPI (grey) is shown. The nuclear signal in the green channel (TAF10 or TAF8 IF) was removed by masking the nucleus and using the “clear” option in ImageJ. Zoom-in regions shown under every image are indicated with a white rectangle. Scale bar (5 μm). **(e)** Boxplot showing enrichment ratios of IF signal at each RNA position over mean cytoplasmic intensity under all the conditions tested. Each grey dot represents one cell. Red horizontal lines are mean values, 95% confidence interval is shown in pink, and standard deviation in blue.

### The position of the interaction domain in the heterodimerization partners defines the co-translational assembly pathways

To further test whether domain position guides co-translational assembly of HFD pairs in TFIID, TAF8 and TAF10 expression vectors were constructed in which the respective HFDs were exchanged. Our nascent RIP experiments from cells co-transfected with these swapped cDNA constructs (*TAF10-HFD8* and *TAF8-HFD10*) resulted in comparable *TAF8-HFD10* mRNA and protein enrichments (Fig. 6a,b and Supplementary Fig. 5a,b); as observed with the corresponding wt constructs (Fig. 2a,b), indicating that the origin of the HFD does not influence the sequential order of co-translational assembly. This experiment also suggested that the position of the HFD (N-or C-terminal), but not its sequence, determines the co-translational pathway by which the protein partners interact. Thus, next we tested whether the co-translational assembly of TAF6-TAF9 HFD pair would follow the simultaneous pathway, as they interact through their N-terminal HFDs (Fig. 6c,d). Our nascent RIPs revealed that both TAF6 and TAF9 co-IP their partners mRNA (Fig. 6c,d and Supplementary Fig. 5c,d), suggesting that they assemble through the simultaneous assembly pathway, presumably as the interaction domains of both proteins are exposed early during their synthesis on the ribosomes. However, we cannot rule out the possibility that the fully synthesized TAF6 or TAF9 could find their respective nascent partners still bound to the ribosomes. Thus, it seems that the position of the dimerization domain may play a critical role in defining the order of co-translational assembly pathway of the corresponding interacting factors.

**Figure 6.**
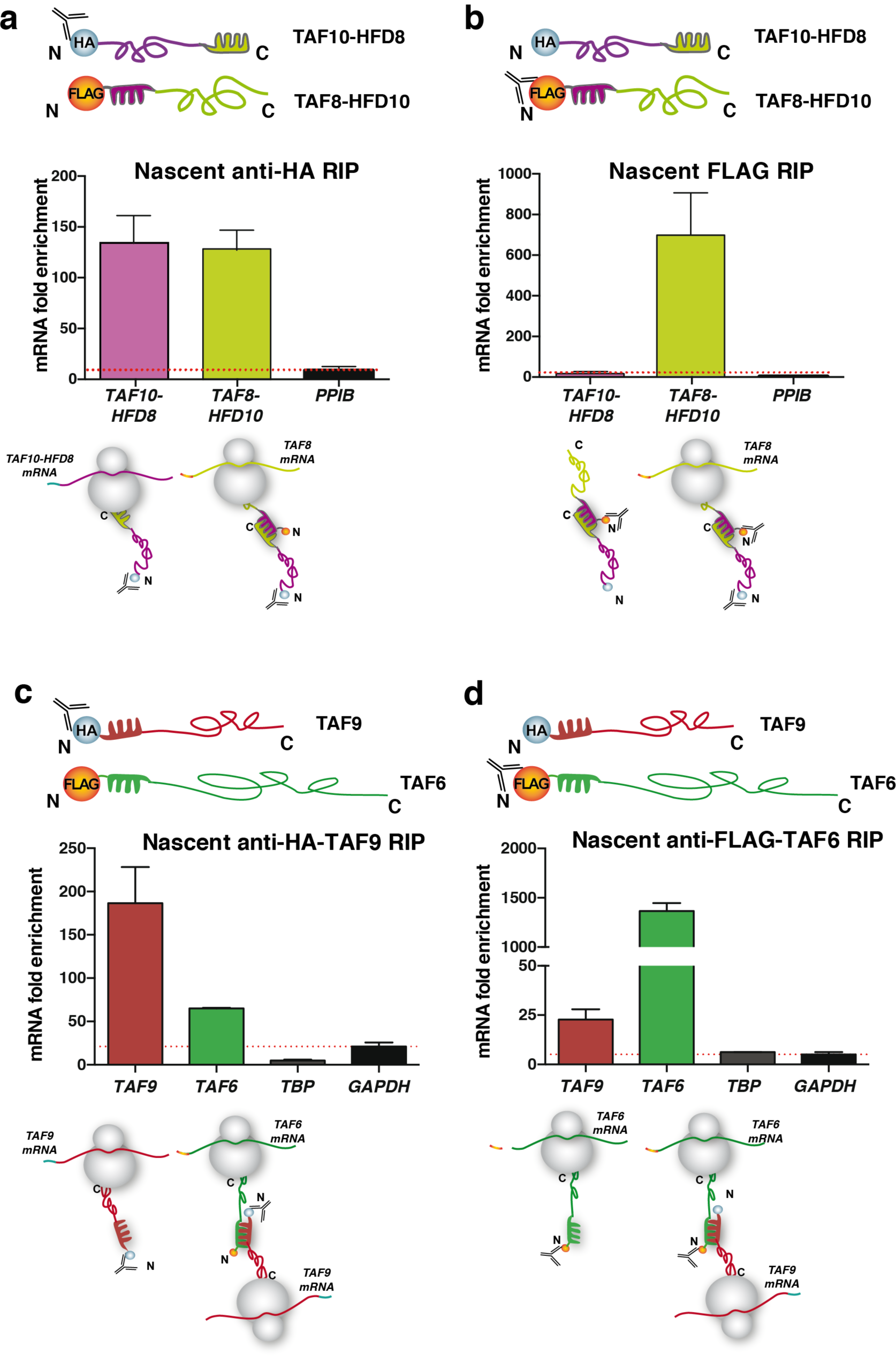
Binding domain position determines the order of co-translational assembly. RIP-qPCR of HFD domain-swapped TAF10 and TAF8 expression constructs using anti-HA **(a)** or anti-FLAG **(b)** antibodies against the respective N-terminal tags. **(c-d)** RIP-qPCR of anti-HA-TAF9 IP (**c**) and anti-FLAG-TAF6 IP (**d**) from HeLa cell polysome extracts cotransfected with TAF9 and TAF6 expression constructs as indicated. *PPIB* **(a-b)** and *GAPDH* **(c-d)** was used as negative control mRNA. In all the graphs error bars represent ±SD from 2 biological replicates.

### The non-HFD-containing TFIID subunits, TBP and TAF1, interact also co-translationally

In TFIID, the evolutionary conserved core domain of TBP interacts with TAF1 via N-terminal TAND region of TAF1 and this interaction modulates the DNA-binding activity of TBP within TFIID^23,24^. To investigate co-translational assembly of other non-HFD-dependent interactions, we carried out genome-wide microarray analysis of TBP-associated RNAs from HeLa cell polysome extracts using a monoclonal antibody against the N-terminus of endogenous human TBP. In addition to TBP mRNA, we detected strong enrichment of 19 coding and non-coding RNAs. Among these, we found mRNAs coding for known TBP-interacting proteins: *BRF1* coding for a factor important for Pol III transcription^25^, *BTAF1* coding for a B-TFIID subunit^26^, as well as *TAF1*, whose enrichment on the microarray was somewhat weaker (Fig. 7a). Nevertheless, RIP-qPCR analysis in human HeLa cells (Fig. 7b) and mouse ESCs (Supplementary Fig. 6a) confirmed the microarray data and revealed a strong enrichment of the *TAF1* mRNA. Quantification of the *TAF1* mRNA in the anti-TBP RIP normalized to the protein IP efficiency indicated that the *TAF1* mRNA enrichment was around 62%. Consistent with the need for active translation, enrichment of all specific mRNAs was lost, or greatly decreased, upon puromycin treatment (Fig. 7b).

**Figure 7.**
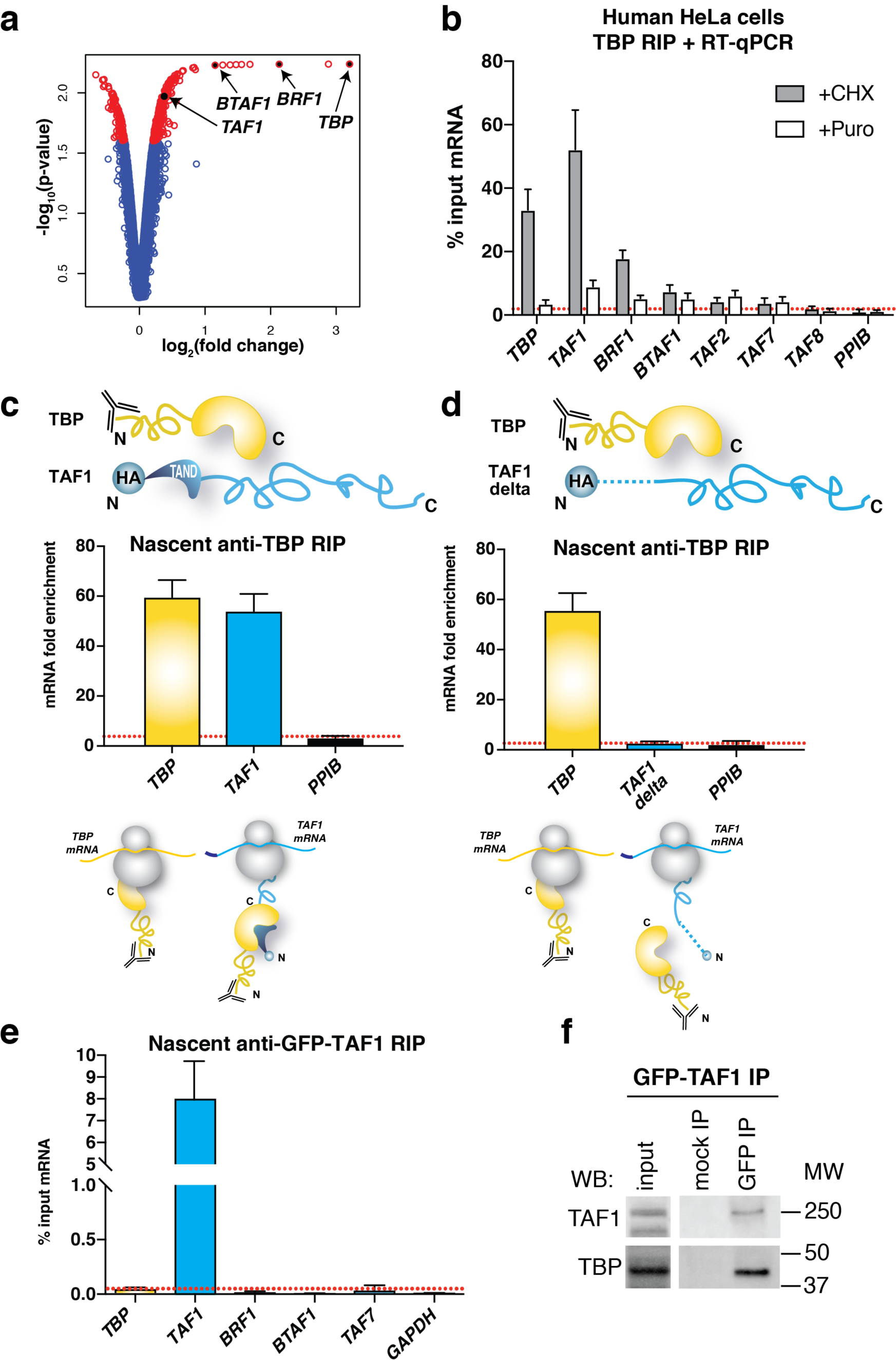
Co-translational assembly of TBP within TFIID and other transcription complexes. **(a)** Microarray results of RIP with an antibody targeting the N-terminus of the endogenous TBP protein. Volcano plot depicting microarray results as log2 of the fold change of IP over mock. A p-value cut-off ≤ 0.025 was applied and corresponding transcripts are in red. Transcripts of interest are highlighted in black. **(b)** RIP-qPCR validation of microarray results in HeLa polysome extracts. **(c-d)** RIP-qPCR using anti-TBP antibody in HeLa cells transfected with TBP and HA-TAF1 expression constructs **(c)** or with TBP and HA-TAF1 with N-terminal deletion of the first 168 amino acids (**d**). **(e)** Anti-GFP RIP-qPCR from polysomes of HeLa cells stably expressing GFP-TAF1. **(f)** Western blot analysis (WB) of GFP IP from polysome extracts prepared from GFP-TAF1 cell line. All error bars are ± S.D. from 3 **(b)** or 2 **(c-e)** biological replicates. Molecular weight (MW) markers are shown in kDa. *PPIB* **(b-d)** and *GAPDH* **(e)** were used as unrelated control mRNAs.

To further investigate the specificity of TBP-TAF1 interaction, we co-transfected expression vectors coding for the full-length human TBP with a ΔTAF1 expression vector, in which sequences coding for the first 168 residues containing the TAND region were deleted. Anti-TBP RIPs from cells expressing ΔTAF1 resulted in complete loss of *TAF1* mRNA enrichment and a reduction of the co-immunoprecipitated protein (Fig. 7c,d, Supplementary Fig. 6b,c). These results are consistent with a requirement of the N-terminal TAF1 domain to recruit TBP to the nascent TAF1 polypeptide. As the protein interface is formed by the C-terminal portion of TBP and the very N-terminus of TAF1^24,27^, we predicted that similarly to TAF8-TAF10 assembly, a sequential assembly is also involved in the TBP-TAF1 interaction. Indeed, nascent anti-TAF1 RIP from an engineered GFP-TAF1 HeLa cell line (Fig. 7e,f) resulted in the enrichment of *TAF1* mRNA, but not that of *TBP*, thus supporting the co-translational assembly of TBP-TAF1 by the sequential pathway.

### TREX-2 and SAGA DUB complexes also assemble co-translationally

To extend our findings beyond TFIID, we examined co-translational assembly of ENY2 subunit with its respective partners. ENY2 is subunit of the TREX-2 mRNA-export complex and the DUB module of the SAGA transcription coactivator^13^. In TREX-2, two ENY2 proteins wrap around the central portion of the large GANP helical scaffold^28^. Similarly, human ENY2 wraps around the N-terminal helix of human ATXN7L3 in the highly intertwined SAGA DUB module^29^ (Fig. 8a). To test whether the co-translational model is generally applicable to multisubunit complexes, we analysed ENY2-associated mRNAs from HeLa cells stably expressing ENY2 with an N-terminal GFP-tag^30^. Interestingly, we found that an anti-GFP-ENY2 RIP co-immunoprecipitates predominantly endogenous *GANP* mRNA and protein (the partner of ENY2 in TREX-2), and also endogenous *ATXN7L3* mRNA and protein (the binding partner of ENY2 in the SAGA DUB module) (Fig. 8b,c). Together, these results demonstrate that co-translational assembly is involved in the assembly of mammalian transcription complexes of diverse architecture and function.

**Figure 8.**
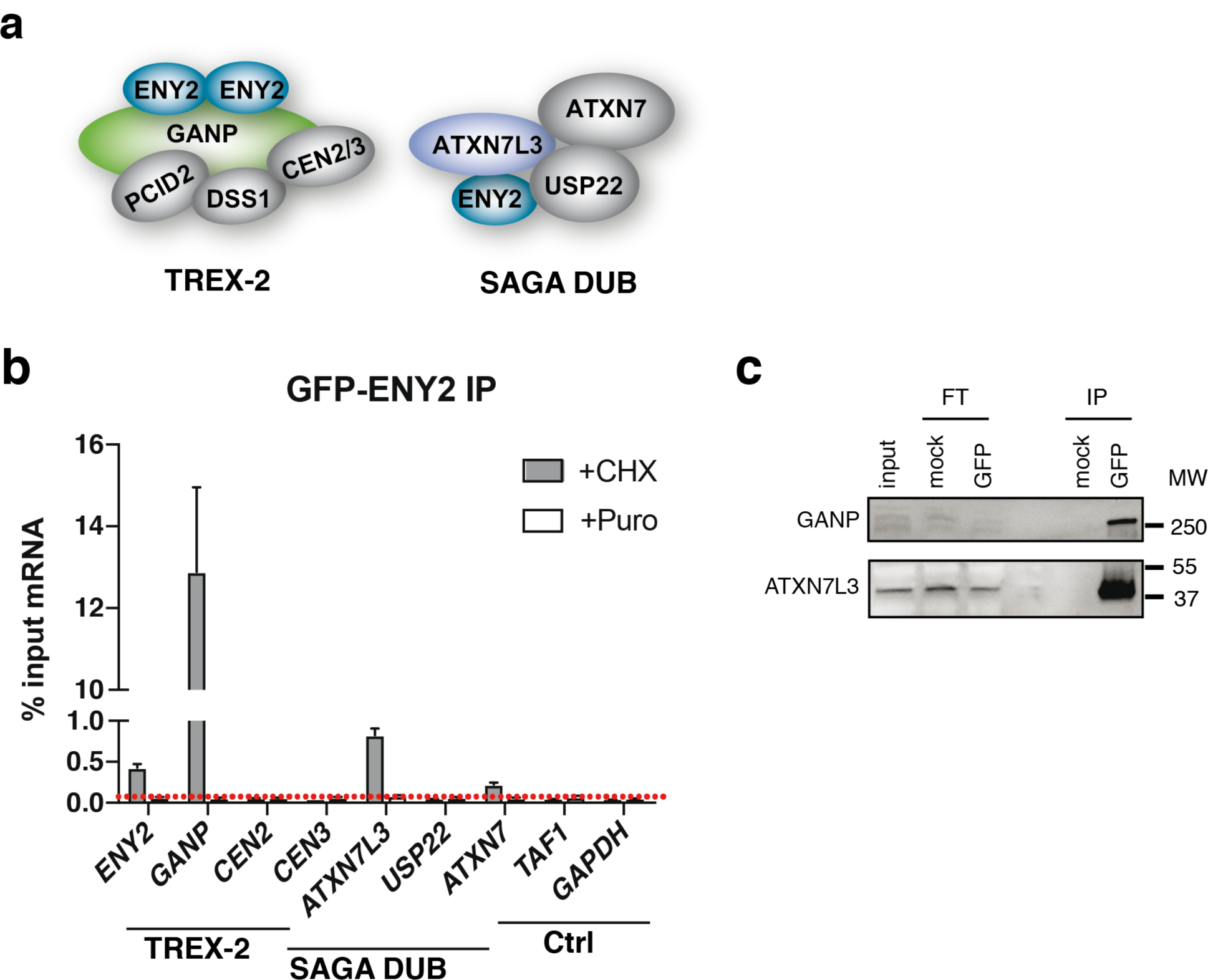
TREX-2 and SAGA DUB complexes also assemble co-translationally. **(a)** ENY2 is shared between TREX-2 and SAGA complex. Two protein molecules of ENY2 binds to the large GANP subunit of TREX-2 complex and ENY2 is also a part of the deubiquitination module of the SAGA complex. **(b)** RIP-qPCR of GFP IP from HeLa cell polysome extracts stably expressing GFP-ENY2 tagged at the N-terminus. Error bars represent ±SD from 2 biological replicates. (**c**) Western blot analysis of GFP IP from polysome extract prepared from GFP-ENY2 cell line. Molecular weight (MW) markers are shown in kDa. FT = flow-through. IP = immunoprecipitation. *GAPDH* was used as unrelated control mRNA.

## Discussion

A functional protein must fold, translocate to its site of action and assemble with the right partners to carry out its function in the cell. The folding and assembly should be a well-regulated process in the cell to avoid non-specific interactions, and also because a single protein might interact with various partners depending on its interaction domain. Most eukaryotic proteins have more than one domain, which enables them to associate with their interaction partners. The building of multi-protein complexes in eukaryotes necessitates co-translational protein folding, the folding of a particular ID while still attached to translating ribosomes, to increase the efficiency of protein synthesis and prevent non-productive interactions^31^. Importantly, co-translational folding is aided by the ribosome, which stabilises specific folding intermediates of a protein^32-34^. Our results further demonstrate that the co-translational dimerization of protein interaction domains directs the assembly of mammalian nuclear multisubunit complexes. The cytoplasmic IF-smiFISH experiments indicate that the described co-translational assembly is clearly occurring in the cytoplasm of human cells and together with the mRNA enrichment calculations show that co-translational assembly is not a minor event. We also show that the position of the heterodimerization domain in a protein could guide its co-translational assembly either by “sequential” or “simultaneous” pathways. These mechanisms could play an important role in maintaining cellular health as excess orphan protein subunits can overburden protein folding and quality control machineries^35^. There is a strong correlation between the amino acid sequence of a protein, its translation rate and co-translational folding^36^. Rare codons in the mRNAs decrease the rate of translation, thereby allowing the protein to fold co-translationally^32^. Interestingly, translation pause sites are located downstream of the ID boundaries in order to regulate proper folding of multi-domain proteins^37^, probably by assuring enough time for the co-translational interaction between the interacting subunits. In good agreement, our KO mESC results suggest that the absence of nascent ID binding with its partner may lead to translational arrest and consequent degradation of both nascent protein and the mRNA coding it. However, further systematic studies need to be carried out in order to study the role of translational pausing in co-translational protein assembly.

Co-translational assembly in homomeric proteins can also cause premature assembly of protein complexes, if two interacting nascent chains are in close proximity. Thus, it seems that homomeric protein IDs are enriched toward the C termini of polypeptide chains across diverse proteomes and it has been suggested that this is the result of evolutionary constraints for folding to occur before assembly^38^. In contrast, our bioinformatics analyses using a limited curated interaction database suggest that in heterodimeric proteins the N-terminal interaction regions are enriched (data not shown), further underlining the idea that co-translational protein assembly in heterodimeric proteins is beneficial for assembling cellular machineries.

The role of chaperones in ribosome associated nascent protein folding is well studied. Hsp70 family of proteins (such as for example yeast Ssb) protect the nascent polypeptide from misfolding and aggregation in eukaryotes^39,40^. In bacteria and yeast, the ribosome-associated chaperones have been shown to interact with the nascent polypeptide chain emerging from the ribosome aiding in its folding^8,41-43^. Moreover, recently it has been suggested that upon emergence of a complete ID the nascent chain interacts with its partner subunit and dissociates the chaperone complex from the nascent chain^8^.

Our results reveal a systemic co-translational building of complexes in mammalian cells, but a thorough proteomic approach is necessary to identify chaperones necessary for these assembly pathways. Note however, that not all mammalian complexes assemble co-translationally, as we could not co-IP any partner mRNAs when carrying out RIPs against two subunits (ZZZ3 and ATAC2) of the human ATAC histone acetyl transferase complex^13^ (data not shown). It is possible that some of the chromatin regulatory complexes assemble through other chaperone-based mechanisms in the cytoplasm or directly in the nucleus.

In summary, we show that building blocks of mammalian nuclear transcription complexes, such as TFIID, SAGA and TREX-2, are assembled during translation and the way in which assembly occurs is consistent with the current knowledge of the preliminary structural organization of the complexes. Moreover, we also studied endogenous TAF10-TAF8 and TBP-TAF1 co-translational assembly in both mouse [mES and NIH3T3 (data not shown)) and human (HeLa and HEK293T (data not shown)] cell lines. Identical results from yeast, mouse and human cells demonstrates that co-translational assembly is a general mechanism in eukaryotes [^8^ and this study]. Thus, the co-translational assembly of multi-protein complexes pathways seems to be a common regulatory mechanism in all eukaryotic cells to ensure efficient solutions to avoid non-specific protein interactions, protein aggregation and probably also to control the correct stoichiometry of subunits belonging to distinct complexes. In addition, our novel findings will significantly advance structural biology studies, because in the future extensive screening experiments will not be required to identify a “real” interaction partner(s) of a given subunit in a multi-protein complex. It will be enough to make an anti-subunit RIP from polysome extracts coupled to microarray analyses (or to RT-qPCRs) and the real endogenous interacting partner(s) can be taken immediately with high confidence for structural determinations and for building the architecture of multi-protein complexes.

## Methods

### Preparation of polysome-containing extracts and RIP

Polysome-containing extracts were prepared from adherent cells harvested at ∼90% confluence by adapting a method for the isolation of ribosomes translating cell type-specific RNAs^44^. Briefly, 10 cm plates were treated with cycloheximide (100 μg/ml final) or puromycin (50 μg/ml final) and returned to the 37^°^C incubator for 15 or 30 minutes, respectively. Subsequently, plates were placed on ice, washed twice with ice-cold PBS and scraped in 500 μl lysis buffer (20 mM HEPES KOH pH 7.5, 150 mM KCl, 10 mM MgCl_2_ and 0.5% (vol/vol) NP-40), supplemented with Complete EDTA-free protease inhibitor cocktail (Roche), 0.5 μM DTT, 40 U/ml RNasin (Promega), and cycloheximide or puromycin as needed. Extracts were prepared by homogenizing cells by 10 strokes of a B-type dounce and centrifugation at 17,000 x g. Clarified extracts were used to start immunoprecipitations, after saving 10% total RNA for input measurement. For TAF10 and TBP IPs, 20 μl of Protein G Dynabeads (ThermoFisher Scientific) were equilibrated by washing 3 times in lysis buffer, resuspended in 400 μl lysis buffer and 2 μl antibody and incubated for 1 hour at room temperature with end-to-end mixing. Beads were washed twice with IP500 buffer (20 mM Tris-HCl, pH 7.5, 150 mM KCl, 10% glycerol (v/v) and 0.1% NP-40 (v/v) and 3 times in lysis buffer. Antibody-bound beads were thus used to perform RIP with polysome extracts overnight at 4 ^°^C with end-over-end mixing. Mock RIP was carried out with equal amount of anti-GST antibody. The next day, beads were washed 4 times for 10 minutes at 4 ^°^C with high salt-containing wash buffer (20 mM HEPES-KOH pH 7.5, 350 mM KCl, 10 mM MgCl2 and 0.1% (vol/vol) NP-40) and subsequently eluted in 350 μl RA1 Lysis buffer and 7 μl 1M DTT. RNAs were purified according to the manufacturer’s instructions of the Macherey-Nagel total RNA purification kit, including the optional on-column DNase digestion step, and eluted twice in the same 60 μl of RNAse-free water. In the case of FLAG, HA, or GFP RIPs, 50 μl packed anti-FLAG M2 affinity gel (Sigma), 50 μl packed EZviewTM Red Protein A affinity gel (Sigma) or 30 μl GFP-TRAP (Chromotek) slurry were equilibrated in lysis buffer and used for RIP.

### cDNA preparation and RT-qPCR

For cDNA synthesis, 5 μl of purified RIP-RNA and 5 μl of 1:10 diluted input RNA samples were used. cDNA was synthesized using random hexamers and SuperScript IV (ThermoFischer Scientific) according to the manufacturer’s instructions. For RIP performed on transfected cells, RNA was additionally treated with Turbo DNase (Ambion) according to the manufacturer’s instructions in order to ensure complete plasmid removal before cDNA synthesis. Quantitative PCR was performed with primers (listed in Supplementary Table 1) on a Roche LightCycler 480 instrument with 45 cycles. In all cases, control cDNAs prepared without reverse transcriptase (−RT) were at least over 10 Cp values of the +RT cDNAs. Enrichment relative to input RNA was calculated using the formula 100*2^[(Cp(Input) − 6.644) - Cp(IP)] and expressed as % input RNA. In the case of RIPs performed on transfected cells, enrichment values were expressed as fold enrichment relative to the mock IP using the formula ΔΔCp [IP/mock], to account for the variability of transient transfections. All experiments were performed with a minimum of 2 biological replicates and values are represented as mean ±S.D. Figures were prepared with GraphPad Prism 6.

### Microarray analysis and library preparation

Polysome extracts and RIP from HeLa cells were performed as described above with mouse monoclonal antibodies 1H8 targeting the N-terminus of TAF10, 3G3 targeting the N terminus of TBP, and 1D10 targeting GST as a nonspecific control (see Supplementary Table 2). Protein G Sepharose beads were used (100 μl beads coupled to 14 μl antibody). After quantification and quality controls performed on Agilent’s Bioanalyzer, biotinylated single strand cDNA targets were prepared, starting from 200 ng of total RNA, using the Ambion WT Expression Kit (Cat # 4411974) and the Affymetrix GeneChip^®^ WT Terminal Labeling Kit (Cat # 900671) according to Affymetrix recommendations. Following fragmentation and end-labeling, 3 μg of cDNAs were hybridized for 16 hours at 45oC on GeneChip^®^ Human Gene 2.0 ST arrays (Affymetrix) interrogating over 40000 RefSeq transcripts and ∼11,000 LncRNAs represented by approximately 27 probes spread across the full length of the transcript. The chips were washed and stained in the GeneChip^®^ Fluidics Station 450 (Affymetrix) and scanned with the GeneChip^®^ Scanner 3000 7G (Affymetrix) at a resolution of 0.7 μm. Raw data (.CEL Intensity files) were extracted from the scanned images using the Affymetrix GeneChip^®^ Command Console (AGCC) version 4.0. CEL files were further processed with Affymetrix Expression Console software version 1.3.1 to calculate probe set signal intensities using Robust Multi-array Average (RMA) algorithms with default settings (Sketch quantile normalization). Statistical analysis was performed using the FCROS package version 1.5.4^45^. Differences are considered significant for p value below 0.025. Volcano plots were performed using RStudio software version 3.3.2. Ribosomal RNA transcripts were filtered out. The microarray results reported in this paper are available in the Gene Expression Omnibus (GEO) under accession number GSE106299.

### Cell lines, cell culture and transfections

HeLa cells grown on adherent plates were obtained from the IGBMC cell culture facility and cultured in a 37^°^C humidified/5% CO2 incubator. Culture media consisted of DMEM supplemented with 1 g/L glucose, 5% Fetal Calf Serum and 40 μg/μl. The GFP-TAF1 cell line was generated by transferring full length human TAF1 fused at its N-terminus to EGFP into HeLa Flp-In/T-REx cells following procedures described in van Nuland et al. (PMID: 23508102). E14 mouse embryonic stem cells (mESCs) at passage 29-31 were obtained from the IGBMC cell culture facility and cultured on gelatinized plates in feeder-free conditions in KnockOut DMEM (Gibco) supplemented with the following: 20 mM L-glutamine, Pen/Strep, 100 μM non-essential amino acids, 100 μM β-mercaptoethanol, N-2 supplement, B-27 supplement, 1000 U/ml LIF (Millipore), 15% ESQ FBS (Gibco), and 2i (3 μM CHIR99021, 1 μM PD0325901, Axon MedChem). Cells were passaged approximately every 3 days. The EGFP-ENY2 cell line was obtained from the IGBMC cell culture facility and maintained as described previously^30^.

Transfections were performed on ∼90% confluent cells in 10 cm plates in antibiotic-free media using Lipofectamine 2000 (Thermo Fisher Scientific) and 3 μg plasmid DNA, according to the manufacturer’s instructions. Medium was replaced with fresh medium containing gentamycin ∼5-6 hours post-transfection and cells were harvested 24 hours later. A descriptive summary of the plasmids used is presented in Supplementary Table 3.

### Protein IP and western blot/antibodies

Antibodies used for RIP, protein IP and western blotting are summarized in Supplementary Table 2. For protein IP, the procedure was performed essentially as for RIP. Bound proteins were eluted in 2 x Laemmli buffer supplemented with 20 mM DTT and boiled 5 min. Subsequently, samples were resolved on SDS-PAGE gels and transferred to nitrocellulose membranes using either wet transfer or BioRad’s Trans-Turbo Blot semi-dry transfer method. Secondary antibodies (goat anti-mouse or rabbit anti-mouse) coupled to HRP (Jackson ImmunoResearch Laboratories) were used at 1:10000 dilution. Signal was revealed using chemiluminescence (Pierce) and detected on the ChemiDoc imaging system (BioRad). For immunoprecipitation using whole cell extracts, 10 confluent 10 cm plates were scraped in PBS containing protease inhibitor (Roche) and resuspended in ∼1 packed cell volume lysis buffer (20 mM Tris-HCl, pH 7.5, 400 mM KCl, 2 mM DTT, 20% glycerol) supplemented with protease inhibitor and 0.5 mM final concentration of DTT. Extracts were prepared by 4 cycles of freezing on liquid nitrogen followed by thawing on ice. The concentration of the clarified extract was measured by Bradford assay and the extract was diluted approximately 1:3 using lysis buffer without salt to achieve a final concentration of ∼150 mM KCl. One mg extract was added to mock-and antibody-bound beads each and IPs were performed as described. Proteins were eluted twice for 5 min at room temperature in 50 μl 0.1 M Glycine, pH 2.8 and neutralized with 3.5 μl 1.8 M Tris-HCl, pH 8.8. 10% of the pooled eluates were resolved on gels.

### Plasmids

The eukaryotic expression plasmid pXJ41 used for all the constructs has been previously described^46^. pXJ41-TAF10-Nter-2HA has been previously described^47^. To generate N-and C-terminally Flag-tagged TAF8, the human TAF8 cDNA was PCR amplified from pACEMam1-CFP-TAF8 (kind gift from Imre Berger, University of Bristol, UK) using primers cotaining EcoR I and Bgl II restriction sites and tags incorporated at the N-or C-terminus, respectively. and digestion by appropriate restriction enzymes. Similarly, C-terminal HA tagged TAF10 was subcloned from pXJ41-TAF10-Nter-2HA by PCR amplification and digestion via restriction enzymes Xho I and Kpn I. The TAF8 mutations, TAF8-HFD and TAF8-HFD-60 amino acids were generated by site-directed mutagenesis using PfuUltra High-Fidelity DNA polymerase (Agilent Technologies), according to the manufacturer’s instructions. The histone fold domain swapped TAF10 and TAF8 constructs were generated by several rounds of PCR amplification using the already mentioned N-terminal tagged TAF10 and TAF8 constructs as template with specific primers and cloned into the vector via restriction enzymes EcoR I and Bgl II. pXJ41-hTBP has been previously described^48^. The HA-TAF1 cDNA^49^ was inserted in pXJ41. TAF1 N-terminal deletion was carried out by site-directed mutagenesis using PfuUltra High-Fidelity DNA polymerase (Agilent Technologies), according to the manufacturer’s instructions. HA tagged TAF9 was subcloned from pSG5-TAF9^50^ by PCR amplification and digestion by restriction enzymes EcoR I and Bgl II. FLAG-tagged TAF6 was also subcloned in a similar manner from pXJ41-TAF6^51^ via restriction enzymes Xho I and Kpn I. All plasmids have been verified by sequencing. Details on the cloning strategies are available upon request. Plasmids are described in Supplementary Table 3.

### Mouse *Taf8* and *Taf10* KO ESC lines

Mice carrying the *Taf8*^*lox*^ allele were bred to mice carrying the *Rosa26*^Cre-ERT2^ allele to produce *Rosa26*^Cre-ERT2/+^;*Taf8*^*flox/flox*^ E3.5 blastocysts and to isolate *Rosa26*^Cre-ERT2/+^; *Taf8*^*flox/flox*^ mouse embryonic stem cells (mESCs)^21^. Similarly, *Rosa26*^Cre-ERT2/R^;*Taf10*^*flox/flox*^ mESCs were derived from *Rosa26*^Cre-ERT2/R^;*Taf10*^*lox/lox*^ E3.5 blastocysts^20^. mESCs were cultured in DMEM (4.5g/l glucose) with 2 mMGlutamax-I, 15% ESQ FBS (Gibco), penicillin, streptomycine, 0.1 mM non-essential amino acids, 0.1% ß-mercaptoethanol, 1,500 U/mL LIF and two inhibitors (2i; 3 μM CHIR99021 and 1 μM PD0325901, Axon MedChem) on gelatin-coated plates. To induce deletion of *Taf8*, mESCs were treated with 0.5**?**μM 4-OH tamoxifen (Sigma) for 5-6 days, and to induce deletion of *Taf10, Rosa26*^Cre-ERT2/R^;*Taf10*^*lox/ lox*^ mESCs were treated for 4 days with 0.1 μM 4-OH tamoxifen (Sigma).

### IF-smiFISH

To visualise proteins and mRNA together, we first performed immunofluorescence (IF) followed by smiFISH. Briefly, cells were treated with 100 μg/ml cycloheximide (Merck) for 15 minutes at 37^°^C, fixed with 4% paraformaldehyde (Electron Microscopy Sciences) for 10 minutes at room temperature (RT), blocked and permeabilized with blocking buffer (10% BSA, 10% Triton-X-100, 200 mM VRC, 2X PBS) for 1 h at 40^°^C, incubated for 2 h at RT with either anti-TAF8 (mouse monoclonal antibody (mAb) 1FR-1B6^52^; diluted 1:1000) or anti-TAF10 (mAb 6TA-2B11^52^; diluted 1:1000) antibody mix followed by incubation (RT, 1 h) with secondary antibody mix Alexa488-labelled goat anti-mouse mAb (Life Technologies, catalogue number A-11001, diluted 1:3000). Following immunofluorescence described above, cells were fixed with 4% paraformaldehyde (Sigma) for 10 minutes at RT. Cells were washed with 1X PBS and incubated with wash buffer [10% Formamide (Sigma) in 2X SSC] for 10 minutes at RT. smiFISH was carried out as previously described^22^. Cells were mounted using Vectashield mounting medium with DAPI (Vector laboratories Inc.). smiFISH primary probes were designed with the R script Oligostan as previously described^22^. Primary probes and secondary probes (Cy3 or digoxigenin conjugated FLAPs) were synthesized and purchased from Integrated DNA Technologies (IDT).

### Imaging and image processing

Confocal imaging of the smiFISH-IF samples was performed on an SP8UV microscope (Leica) equipped with a 633-nm HeNe laser, a 561-nm DPSS laser, a 488-nm argon laser and a 405-nm laser diode. A 63x oil immersion objective (NA 1.4) was used and images were taken by using the hybrid detector photon-counting mode. The z-stacks were taken with a z-spacing of 300 nm for a total of 4-6 μm. Image processing was performed using the Fiji/Image J software. All images were processed the same way. In detail, one cell of an image was cropped and one representing z-slice per cell was chosen. Additionally, one single IF or smiFISH spot from the corresponding cells was cropped as well. Afterwards, the channels of the different images were split and grey values were adjusted to better visualize the spots in the cytoplasm. The nuclear signal in the green channel (TAF10 or TAF8 IF) was removed by masking the nucleus and using the “clear” option. Finally, the processed channels were merged again.

### Image analysis and data presentation

To measure the degree of spatial overlap of smiFISH (mRNA) and IF (protein) signal, an enrichment ratio was calculated as described next. Such a quantification was opted in order to take into account the variability of IF signal between cells, making single object detection in this channel difficult. Cells and nuclei were outlined manually in 2D based on the GFP and DAPI image, respectively. Subsequent analyses were restricted to the cytoplasm. mRNAs were detected in 3D with FISH-quant^22^. Identical detection settings were used when different experimental conditions were compared for the same gene. Each cell was post-processed separately. First, the median pixel intensity in the IF image at the identified RNA positions was calculated. Second, a normalization factor as the median IF intensity of the outlined cytoplasm within the z-range of the detected mRNAs was estimated. The enrichment ratio of the cell was then calculated as the ratio of the median IF intensity at the RNA positions divided by the mean cytoplasmic intensity. Boxplots of enrichment ratios were generated with the Matlab function notBoxPlot. Each dot corresponds to the estimation of one cell. Horizontal lines are mean values, 95% confidence interval is shown in red, and standard deviation in blue. Statistical comparison between different experimental conditions were performed with two-sample Kolmogorov-Smirnov test (Matlab function kstest2).

## Acknowledgements

We would like to thank all members of the Tora lab for thoughtful discussions and suggestions throughout the course of the work, and G. Caliskan for help with the ATAC experiments. In addition, we are grateful to E. Bertrand and his group for smiRNA FISH training, T. Sexton for advice and carefully reading the manuscript, V. Alunni for microarray sample preparation, S. Bour for illustrations, K. Gupta and I. Berger for suggesting TAF deletions and mutations, D. Van Essen for providing constructs, P. Jane Palli and G. Travé for help with protein domain analyses, the IGBMC cell culture facility for cells and media.

## Funding

This work was supported by funds from CNRS, INSERM, and Strasbourg University. This study was also supported by the European Research Council (ERC) Advanced grant (ERC-2013-340551, Birtoaction, to LT) and grant ANR-10-LABX-0030-INRT (to LT), a French State fund managed by the Agence Nationale de la Recherche under the frame program Investissements d’Avenir ANR-10-IDEX-0002-02 and a grant from the Collaborative Center for X-linked Dystonia and Parkinsonism (to HTMT).

## Authors’ contributions

IK, PM and LT designed the study; IK and PM performed all the molecular lab work, JMG performed all the cloning experiments. PM and SConic carried out the imaging experiments, which were analysed by FM, SCaponi created a stable cell line, FES, PB and SDV carried out the mouse ESC knock-out experiments; IK, PM analysed data, DD analysed the microarray data. All authors contributed to the text and figure panels. IK, PM, HTMT, SDV and LT wrote the manuscript. All authors gave final approval for publication.

## Competing Interests

The authors declare that they have no competing interests.

